# Visually driven neuropil activity and information encoding in mouse area V1

**DOI:** 10.1101/113019

**Authors:** Sangkyun Lee, Jochen F. Meyer, Stelios M. Smirnakis

**Affiliations:** Department of Neurology, Brigham and Women’s Hospital, Jamaica Plain Veterans Administration Hospital, Harvard Medical School, Boston, MA, 02115, USA; Department of Neurology, Baylor College of Medicine, Houston, Texas, 77030, USA

## Abstract

Spontaneous calcium fluorescence recorded from large cortical neuropil patches strongly correlates with the electro-corticogram, and is thought to arguably reflect primarily pre-synaptic inputs. Here we used in vivo 2-photon imaging with Oregon Green Bapta (OGB) to study neuropil visual responses to moving gratings in layer 2/3 of mouse area V1. We found neuropil responses to be more reliable and more strongly modulated than neighboring somatic activity. Furthermore, stimulus independent modulations in neuropil activity, i.e. noise correlations, were highly coherent across the cortical surface, up to distances of at least 200 μm. Pairwise neuropil-to-neuropil-patch noise correlation strength was much higher than cell-to-cell noise correlation strength and depended strongly on brain state, decreasing in quiet wakefulness relative to light anesthesia. The profile of neuropil noise correlation strength decreased gently with distance, dropping by ~12% at a distance of 200 μm. This was comparatively slower than the profile of cell-to-cell noise correlations, which dropped by ~30% at 200 μm. Interestingly, in spite of the “salt & pepper” organization of orientation and direction encoding across mouse V1 neurons, populations of neuropil patches, even of moderately large size (radius ~100μm), showed high accuracy for discriminating perpendicularly moving gratings commensurate to the accuracy of corresponding cell populations. These observations underscore the dynamic nature of the functional organization of neuropil activity.

**Conflict of Interest:** The authors declare that no competing interests exist.

## Introduction

Two-photon microscopy allows imaging of both cellular and neuropil activity [1–5]. Chklovskii et al. [6] and Braitenberg et al. [7] argued that axons (including axonal boutons) and dendrites constitute ~70-80% of the neuropil. Neuropil activity reflects chiefly aggregate activity from axonal and dendritic branches as well as pre- and post-synaptic components in large numbers of local synaptic aggregates. Kerr and colleagues measured the spontaneous calcium signal modulation in large ~10,000 μm^2^ patches of neuropil in layer 2/3 of mouse motor and barrel cortices [2] and showed that it correlates well with the simultaneously recorded electro-corticogram (ECoG). In fact, Kerr et al. further argued that neuropil calcium signal modulations mainly reflect presynaptic inputs, since blocking post-synaptic transmission did not alter neuropil calcium signal amplitude [2]. Studying the spatial organization and properties of visually driven neuropil activity is of interest, as it may provide clues about the emergence of adjacent neuronal somatic activity and neuronal population interactions. For example, we do not know the degree to which aggregate neuropil activity in the vicinity of neurons retains orientation/direction selectivity bias, or the relation between the reliability of neuropil visual responses and the reliability of adjacent neuronal responses, as well as their relation to brain state. In what follows we explore some of these questions in L2/3 of mouse primary visual cortex.

Sensory processing in the brain has been extensively studied under anesthesia, and it is true that several functional properties of neurons remain unchanged when measured in the anesthetized versus the awake state [8]. However, neuronal activity is, in general, brain-state dependent. For instance, under light anesthesia, visually evoked responses are stronger than responses measured in the quiet awake state, and there is an increase in population synchrony [9,10]. This raises the question whether changes in neuronal activity and synchrony as a function of brain state have a correlate in corresponding neuropil activity changes.

We recorded neuronal activity using *in vivo* two-photon calcium imaging in layer 2/3 of mouse primary visual cortex (V1) while presenting drifting grating stimuli subtending a large visual angle. Our experiments reveal that local neuropil patches exhibit stronger and more reliable calcium responses to visual stimulation than adjacent neurons, and this difference is more pronounced under anesthesia than during quiet wakefulness. Neuropil activity is highly correlated across the field of view but correlation strength decays slowly as a function of distance up to the distance examined (~200 μm). Neuropil correlation strength depends on brain state, being higher under light anesthesia compared to quiet wakefulness. Finally, somewhat surprisingly because of the “salt & pepper” mouse V1 organization, relatively large (~15×15 μm^2^ or larger) neuropil patches show high decoding accuracies in a direction discrimination paradigm, on par with the performance of nearby cell populations. This suggests that in layer 2/3 of mouse V1, substantial local direction information is contained in the aggregate activity of neuropil patches with radii ranging from 30 to as large as 200 μm.

## Materials and Methods

### Animal preparation

All experiments and animal procedures were performed in accordance with guidelines of the National Institutes of Health for the care and use of laboratory animals and were approved by the IACUC at Baylor College of Medicine. All mice used were derived from C57BL/6 lines and were 4 to 8 weeks old. Experiments under anesthesia were performed in 5 fields of view (FOV’s) from 3 Parvalbumin (PV)-Cre X Ai9 F1 mice and 2 FOV’s from 2 Dlx5/6-Cre X Ai9 F1 mice. Awake experiments were performed in 11 FOV’s (2 FOV’s from 2 PV-Cre X Ai9 F1 mice and 9 FOV’s from 4 wild-type C57BL6 mice).

### Surgery

All procedures were carried out according to animal welfare guidelines authorized by the Baylor College of Medicine IACUC committee. All surgeries were performed under general anesthesia with 1.5% isoflurane. The mouse head was fixed in a stereotactical stage (Kopf Instruments), and eyes were protected with a thin layer of polydimethylsiloxane (30,000 cst, Sigma-Aldrich). After removing the scalp, a custom-made titanium headplate was attached to the skull with dental acrylic (Lang Dental). A 3 mm wide circular craniotomy centered 2.5 mm lateral of the midline and 1.2 mm anterior of the lambda suture was made, targeting the middle of the monocular region of left V1. A coverglass with a hole for pipette access was placed on the brain and carefully anchored with vetbond glue (3M, Saint Paul, MN) and dental acrylic (Lang Dental).

### Dye loading and imaging

*We used the calcium indicator Oregon Green BAPTA-1 (OGB) because it stains uniformly both cell bodies and aggregate neuropil processes near the site of injection*. 50 μg Oregon Green 488 BAPTA-1 AM (OGB, Invitrogen) was dissolved in 4 μΙ DMSO (heated to 40^0^ C) with 10% Pluronic acid F-127 (Invitrogen), vortexed for 20 min, and diluted in 40 μl 0.9%-NaCl solution containing 10 μΜ Alexa-594 for experiments with tdTomato-labeled interneurons, and 10 μΜ Sulforhodamine 101 [11] for selective astrocyte-labeling in other experiments. This solution was injected using a glass pipette at depths of 200, 300, and 400 μm of mouse visual cortex under two-photon visual guidance. Cell imaging commenced 1 hour after the dye injection. Populations of 50-100 cells located 150-250 μm below the pia were imaged with water-dipping objective lenses, either 20x, 0.95 NA (Olympus), or 25x, 1.1 NA (Nikon), in a modified Prairie Ultima IV two-photon laser scanning microscope (Prairie Technologies, Middleton, WI), fed by a Chameleon Ultra II laser (Coherent, Santa Clara, CA). Local windows of 200250 μm × 200-250 μm with an in-plane iso-symmetric pixel resolution of 1.2 – 1.9 μm were imaged at frame rates of 7-10 Hz. Depending on imaging depth, the laser power was kept between 10 mW at the surface and 50 mW at depths below 250 μm, at 840 nm (when the patch pipette was filled with Alexa 594 or sulforhodamine) or 890 nm (when filled with dextran) laser wavelength. During visual stimulation experiments under light anesthesia, 0.7% isoflurane was maintained during the experiment via a nose cone. The body temperature of the mouse was kept at 36-37^0^ C with a heating pad (Harvard Apparatus).

### Patch-clamp recording

Whole-cell and loose-patch recordings were obtained with a Heka EPC-10 USB amplifier in current-clamp mode using standard techniques [12]. Glass pipettes of 6-8 MOhm, filled with intracellular solution (in mM: 105 K-gluconate, 30 KCl, 10 HEPES, 10 phosphocreatine, 4 ATPMg, and 0.3 GTP), adjusted to 290 mOsm and pH 7.3 with KOH [13] and containing 10 μΜ Alexa-594 or tetramethylrhodamine dextran (Invitrogen), were used for the recording under two-photon visual guidance. After approaching a target cell with pressure adjustment based on the depth of the pipette from the pia and the distance to the cell, a GigaOhm seal between cell membrane and the pipette was formed. A patch of cell membrane was broken by applying 200 ms pulses of negative pressure with increasing strength using a picospritzer III (Parker Hannifin, Pine Brook, NJ). Fast pipette capacitance was neutralized before break-in, and slow capacitance afterwards. We targeted pyramidal cells in layer 2/3 (between 100 and 250 μm below the pia).

### Visual stimulation

Visual stimuli were generated in MATLAB and displayed using Psychtoolbox [14]. The stimuli were presented on an LCD monitor (DELL 2408WFP, Dell, Texas, USA) at 60 Hz frame rate, positioned 32 cm in front of the right eye, centered at 45 degrees clockwise from the mouse’s body axis. The visual angle of the screen spanned 54^0^ elevation and 78^0^ azimuth. The screen was gamma-corrected, and the mean luminance level was photopic at 80 cd/m^2^. Our visual stimulation paradigm consisted of grating stimuli moving in one of 2 orthogonal directions (0^0^ vs 90^0^) at 3 different contrast levels (100%, 40%, 15% Michelson contrast [15]). Each grating stimulus was generated as a square wave with a spatial frequency of 0.04 cycles/degree and a temporal frequency of 0.5 Hz. Grating stimuli were presented for 600 ms (500 ms for a single FOV) followed by an inter-stimulus interval of 1.5 seconds during which a full-field gray screen at the same mean luminance (background illumination) was presented. Each of the 6 possible conditions was presented 80-100 times in pseudo-random interleaved order.

## Data analysis

### Preprocessing

Movies were motion-corrected along a 2D image plane (x-y motion). Motion parameters were estimated in the red channel, in which tdTomato-labeled interneurons were identified, by registering all image frames to the average of the first 5 image frames using a sub-pixel registration method [16]. Then, the correction parameters were applied to the green channel (in which calcium dynamics of cells were monitored) to reconstruct motion-corrected movies. Data from awake animals were further constrained based on the extent of their movements during imaging. All trials showing any image frames with movement >2 × pixel size (~3μm within the x-y plane) from the first frame of each movie were excluded in the data analysis. The resulting motion level was similar to the x-y motion level of anesthetized animals. The application of this criterion reduced the number of trials per condition to ~70 but minimized the possibility that different results between the different brain states were due to motion artifacts.

For cell identification, a local region-of-interest (ROI) over a cell body was manually defined with a circular disk to cover the cell body, then scanned for the pixel with the highest fluorescence value within the disk (cell body center). The boundary of the region containing the cell signal was then defined by thresholding at 0.5 × maximum fluorescence within the disk along the polar coordinates (Figure 1A) [17]. To correct for slow signal drift over time, the signal time series from each pixel was high-pass filtered (HPF) at 0.05 Hz using the discrete cosine transformation. We then measured the ratio between the mean calcium signal within the lumen of non-radial blood vessels (≤10 μm) and the surrounding neuropil patch [4]. This gave us an approximate measure of neuropil contamination at the cell soma, the so-called contamination scale, whose typical value was 0.6. To correct for the neuropil contamination at the soma, the mean fluorescence of the adjacent neuropil patch, F_n_, was subtracted using the contamination scale S: F_correct_ = F - S*F_n_. The patch F_n_ formed an annulus with a radius of 7-15 μm centered around the soma, excluding pixels that belonged to other cell bodies or astrocytes, which are labeled red with sulforhodamine (Figure 1A).

**Figure 1.**
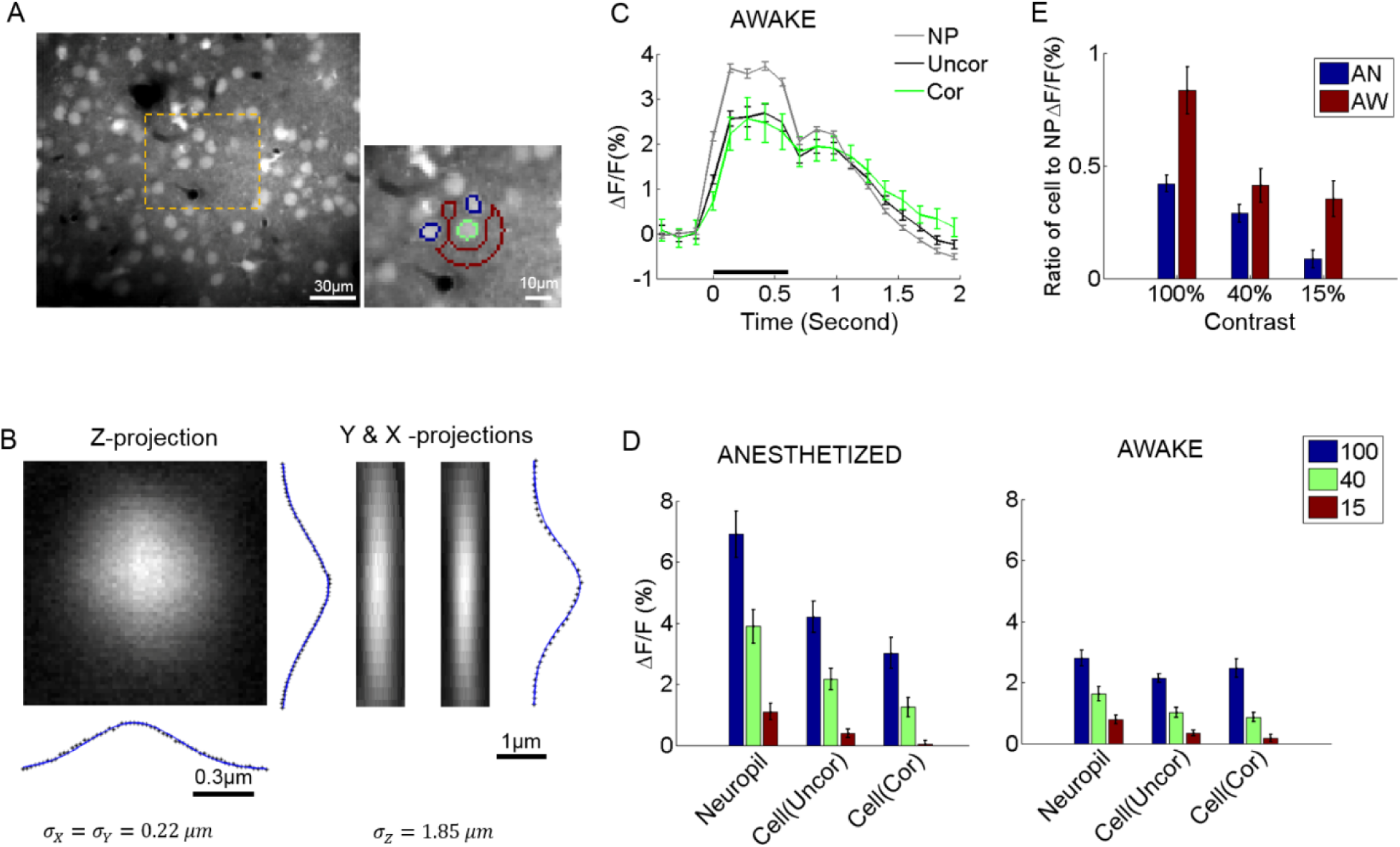
Neuropil and somatic response variability and dependence to contrast. **(A)**Selecting cell somata and local neuropil patches. An example of a typical field of view (FOV, left). The orange area is magnified on the right. Cell radii in our experiments were generally <=5 μm. A local neuropil patch was defined around each cell body as an annulus with inner radius 7 μm and outer radius 15 μm. Any annulus pixels belonging to or being close to other cells (i.e., within 7 μm of the cell center), vessels, or poorly stained regions were excluded by visual inspection to minimize contamination of the neuropil signal by sources other than neuropil. In the example, the green outline shows a selected cell soma (green outline only for visualization). Its local neuropil patch is outlined in red. Note that nearby neurons were excluded from the definition of the neuropil ROI (annulus) and are shown in blue (outline only). The pixel resolution is 1.2×1.2 μm. (B) Gaussian fit of the point spread function (PSF) of our microscope measured by imaging a single spherical fluorophore with a radius of 0.1 μm. (C) Example of the average response of cells and neuropil-patches (annular patch with radii of 7 to15 μm) in a single FOV to 100% contrast. “NP”: mean neuropil response. “Uncor”: mean uncorrected somatic response. “Cor”: mean somatic response after neuropil contamination correction (see Methods) (D) Mean evoked fluorescence responses to gratings of 100%, 40%, and 15% contrast, respectively, derived from all visually responsive cells in 7 anesthetized (AN) and 11 awake (AW) FOV’s, respectively. (E) Ratio of mean somatic versus mean neuropil responses (pooled from all FOV data). These ratios are plotted after correcting for neuropil contamination. Note that the relative strength of cell versus neuropil ΔF/F responses decreased in the lightly anesthetized state. This is largely due to the fact that neuropil responses markedly increase in the lightly anesthetized state. NP: neuropil, AW: awake state, AN: anesthetized state. Statistics are across FOV’s.

In our calcium imaging setup, the point spread function is wider along the z-axis (Figure 1B), so the correction factor mostly compensates for neuropil contamination along the z-axis. The contamination of the neuronal signal by the neuropil signal varied with cell size as well as with the in-plane diameter of the soma cross-section. We therefore examined a range of values around the empirically estimated value S = 0.5 - 0.6 (Supplementary Figure 2), and verified that reasonable variation in the level of contamination (S) did not significantly affect our conclusions (Figures 1C-D and Supplementary Figure 1). We found that a range of correction factors S (S = 0.4 - 0.7) result in similar response patterns as shown in Figures 1C-D. Higher levels of contamination (S = 0.8) began to distort visually evoked responses, resulting in prolonged, non-physiological, delays of the visual response peak (Supplementary Figure 1 A-B, D-E). Such high levels of contamination were not empirically found in our data, as judged by measuring the spread of the calcium signal into the lumen of vessels running parallel or perpendicular to the field of view, whose lumens were commensurate to the typical range of cell sizes.

#### Effects of Correction on noise correlations

The neuropil correction naturally decreases somewhat the magnitude of the cell ΔF/F response as it removes the component of it that is due to neuropil contamination (Figure 1, Supplementary Figure 1). This decrease is mainly evident in the anesthetized state, where neuropil response is relatively higher. Its main effect however, is in allowing a veridical estimation of noise correlation strength. Prior to neuropil contamination correction, mean inter-neuronal noise correlation coefficients ranged from ~0.15 to 0.3 (Supplementary Figure 2), commensurate with results from several published 2-photon imaging studies (Kerr JN et al. 2007; Golshani P et al. 2009; Rothschild G et al. 2010). Such measurements are subject to neuropil contamination, which, because of its high spatial coherence, has the potential to substantially alter the magnitude of noise correlations. Neuropil contamination correction results in a significant adjustment resulting in noise correlation coefficients ~ 0.05 (Supplementary Figure 2). Note that we calculated the noise correlation by estimating the spike rates from the corrected neuropil signal (see below for spike-rate estimation). These values are much closer to values reported in recent electrophysiology studies (Ecker AS et al. 2010; Ecker AS et al. 2014), which had excellent single unit isolation and recording stability. In conclusion, while the correction for neuropil contamination has little effect on mean neuronal visual responses, it does have a big effect on the strength of noise correlations across pairs of cells.

*Note that unless explicitly mentioned, all cell responses are calculated by implementing a neuropil contamination correction with the empirically determined factor S = 0.6*.

Only putative excitatory pyramidal neurons and their surrounding neuropil patches were used in the following analyses. Interneurons expressing Td-tomato and astrocytes (stained red with sulforhodamine) were excluded from the analysis.

### A New, Stable, Spike-Estimation Algorithm Outperforming [18]

We used a new method based on sparse non-negative linear regression to estimate spike rates associated with the calcium fluorescence ΔF/F signal. This method assumes linear calcium dynamics with a time constant that does not change over the course of the experiment. Cell firing is modeled as causing an instant (within one ~130 ms frame) calcium increase that slowly decays in the subsequent frames:

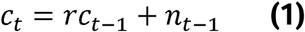

where *c* is the cell’s calcium Δ F/F signal, t indexes time in units of ~130 ms frames, *r* = 1 -Δ/τ, where Δ is the frame duration, τ the time constant and n the normalized spike number during the frame duration, which has the same units as ΔF/F.

In Matrix-vector form,

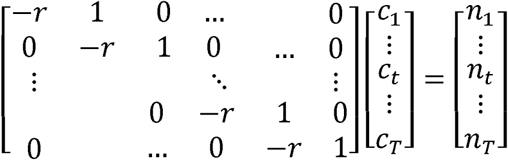

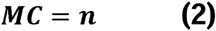

where ***M*** is a convolution matrix that transforms a calcium concentration time series to spikes, and *n* > 0.

Or

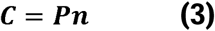

where ***P*** is the inverse matrix of ***M***, and ***n*** > 0.

Assuming exponential distribution of spikes, the objective function for ***n*** is defined as a minimum mean square error form:

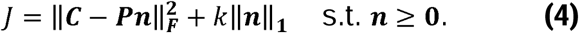

where *k* is a regularization parameter.

After incorporating a term (***a*** > **0**) that allows us to optimize the spatial filtering of the pixels within the cell body, this formula becomes:

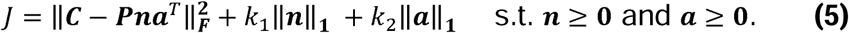

*C* is now a multi-pixel Matrix (time samples x pixels), and the spatial filter *a* is constrained to be non-negative and to be bounded for convergence of the objective function. Here we adopted the L_1_-norm (| |_i_) minimization to bound *a* and *n* guaranteeing the convergence of the alternating optimization algorithm (between n and a). The advantage of the regularization used for *a* is to minimize contributions of low SNR pixels to spike estimation.

To estimate *n* and *a* iteratively, we used an optimization method similar to the Expectation Maximization algorithm [19], alternately estimating *n* while holding ***a*** fixed and vice versa:

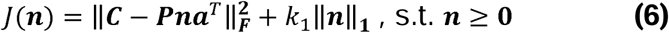

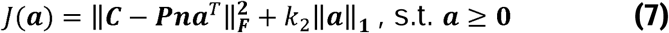

This model can also be interpreted as a Bayesian model, maximizing the a posterior probability

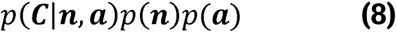

where

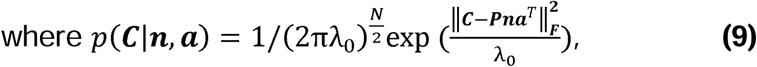

with priors

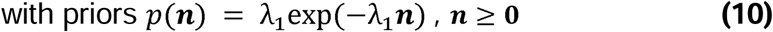

and

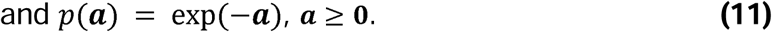

From the Bayesian model, the parameter k_1_ and k_2_ can be released freely by maximizing the posterior probability. Specifically, this was performed by alternatively updating ***C***, ***n***, and ***a*** and updating only λ_0_ with λ_1_ fixed. In our estimation, λ_1_ was set as the imaging frame period. Therefore, for Eqs. (6) and (7), both parameters, ***k_t_*** and ***k_2_*,** can be controlled solely with λ_0_.

In each iteration, after estimating ***n*** and ***a***, λο can be updated by maximizing

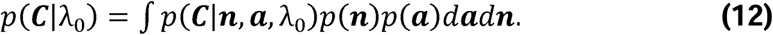

However, due to the intractability of the integral, it is a common practice to obtain the λ_0_ estimate by setting the derivative of the log-posterior probability, log(*p*(***C***|***n***, ***a***)*p*(***n***)*p*(***a***)), with respect to equal to 0 and then solving the equation.

We used an optimization technique for the linear regression model with the L_1_ norm, which uses a log-barrier technique for non-negative constraints and tests for convergence of the learning algorithm via assessing the gap between primal and dual problems [20,21]. This convex optimization with completely bounding alternative parameters theoretically converges to the global minimum [22,23].

Our spike estimation method, like a previously published method [18], assumes linear calcium dynamics and an exponential distribution of spike rates, and uses alternating optimization and the log-barrier optimization technique for L_1_-norm minimization. However, it has one major advantage over [18], which is that its parameter optimization is stable. Theoretically, interacting parameters could cause the objective (cost) function to become unstable because the same objective value can result from a different combination of parameter values. For instance, the parameters σ and λ for exp(-(…)^2^/ σ)exp(-λ(…)) in Eq.11 of the Vogelstein algorithm [18] are interacting and can lead to unstable optimization (Figure 2). However, our method does not have this problem, because the sub-objective functions, Eqs. (6) and (7) have only one single free parameter each, *k_t_* and *k_2_* respectively, as they are optimized alternatingly with respect to ***n*** and ***a***. Both *k_t_* and *k_2_* are determined solely by λ_0_.

**Figure 2.**
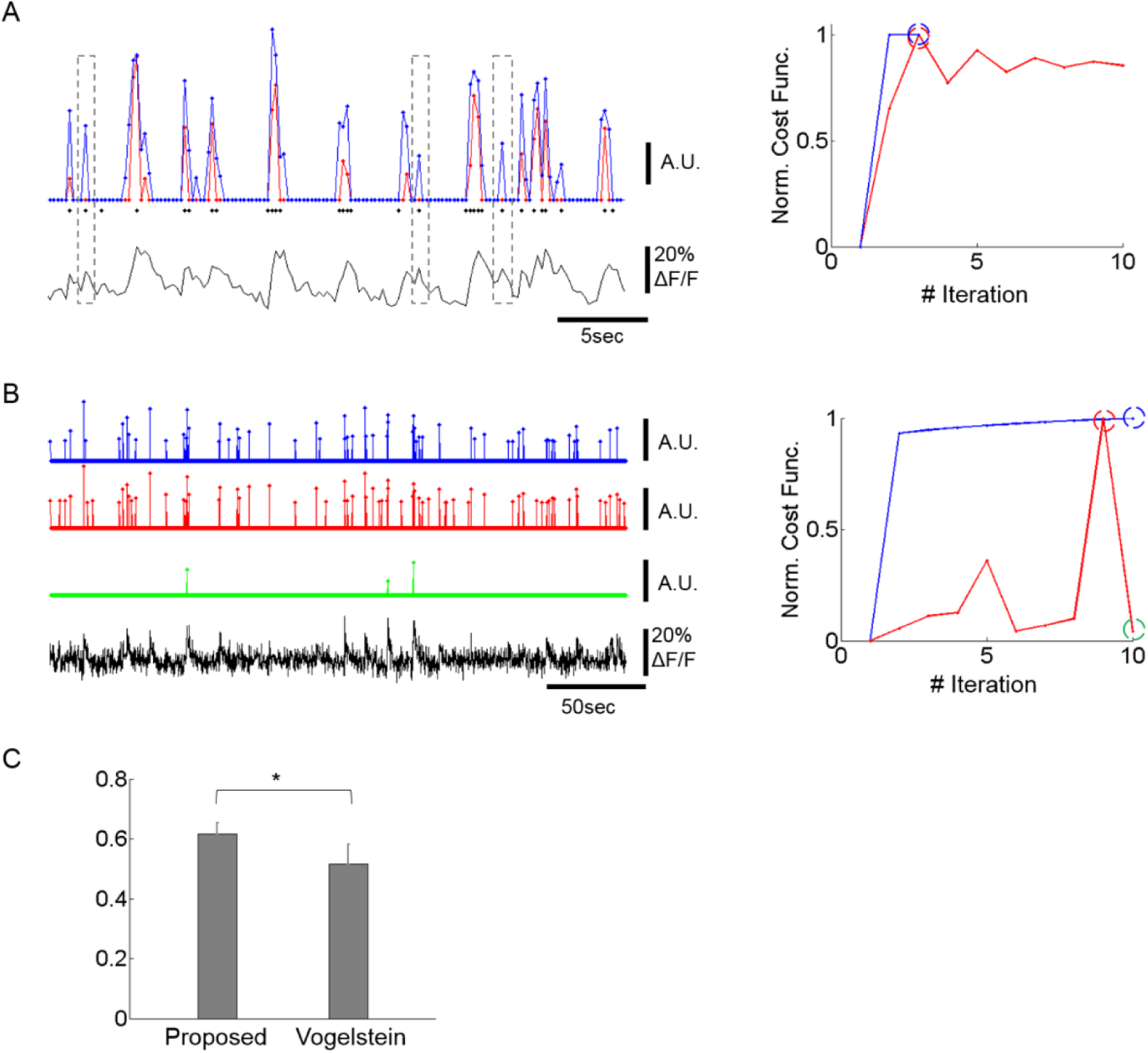
Improving spike inference from the calcium signal. **(A)** Example of spike trains inferred by our deconvolution algorithm and by a previously published method (Vogelstein JT *et al.* 2010). *Left bottom:* OGB calcium fluorescence trace of a cell whose spikes were measured via patch-clamp recording. *Left Top:* The prediction of our deconvolution method (blue) versus the Vogelstein method (red, Vogelstein JT *et al.* 2010). Actual spike events at the time of image frames are overlaid as black dots. The events do not include the spike numbers. For major calcium transients both methods show similar predictions. For intermediate and low calcium transients, however, our method identifies action potentials with greater sensitivity than the Vogelstein method (see dotted outlines). This is reflected in a higher correlation of the deconvolved to the actual spike rates (0.64 for ours, 0.51 for Vogelstein). *Right:* Convergence of our method (blue) versus Vogelstein’s method (red) as a function of the number of iterations, applied to the time-series of a cell’s calcium signal. Blue and red dashed circles indicate the iteration at which algorithms return the optimal estimates. For visualization, each estimated spike-rate train was thresholded at 3x standard deviation. Since spike trains are sparse, thresholding did not significantly suppress spike rates. In particular, the green trace was not affected by this thresholding process. **(B)** Example of spike trains inferred by the two methods in a typical calcium signal recording. *Left:* Both methods identify strong calcium transients (blue and red) at the maximum of the cost function within 10 iterations performed. The green graph shows spike identification at an iteration (iteration = 10), where the spike identification is markedly poorer than in iteration nine, where the best result was obtained. The same thresholding for visualization as (A) was applied. *Right:* Learning curve of the two methods. The blue, red, and green dashed circles indicate the iterations at which the estimates were obtained. While the learning curve of the Vogelstein method does not converge after multiple iterations, the proposed method shows a monotonic increase of the learning curve. This illustrates that the Vogelstein method can be inherently unstable, whereas the method proposed here is stable by converging monotonically. **(C)** Correlation between deconvolution algorithms and actual spike trains for 5 patched and imaged units. The proposed method shows significantly higher correlation (p<0.05; paired t-test, n=5) than the Vogelstein algorithm. * p<0.05.

The same instability occurs with respect to estimating the spatial filter, ***α***, and the estimated calcium trace, **C** (Eqs. 33 and 36 of [18]). Moreover, the spatial filter ***α*** in the

Vogelstein algorithm is unconstrained and could be negative thus potentially yielding erroneous estimates. In contrast, our method constrains the spatial filter **a** to be both non-negative and upper-bounded. This prevents the main objective function from oscillating through alternating optimizations of ***n*** and ***a*.**

Application of our method reliably estimated the actual spike rate (Figure 2A left and C) and outperformed the Vogelstein algorithm by producing higher correlation (25% higher, p<0.05) to actual spikes simultaneously recorded (Figure 2C). Data from an optical-imaging only session also showed reliable results by identifying rapid calcium transients as spikes (Figure 2B). In addition, our method shows clear empirical convergence of the learning curves (e.g., Figure 2A and B) together with the above theoretical guarantee in accordance with convex optimization theory.

### Cell and Neuropil Analysis

To estimate the spike rate, the pre-processed (HPF) fluorescence signal was normalized, pixel by pixel, by calculating (F-F_0_)/F_0_ (i.e., ΔF/F). F_0_ was defined as the mean of the fluorescence values that were less than 2 standard deviations above the overall mean for that pixel. This value was chosen to exclude outlier values, which were likely related to spike activity. Spike rates were then estimated by applying the method described above (Figure 2) to the pixels constituting a cell body. Following spike train estimation, a threshold of 1/2 × standard deviation of the spike rate was used to suppress spurious spikes arising from photon noise. This threshold was found to work well in that it was less likely to suppress activity generated by actual spikes, because spike-rates were sparse and exponentially distributed and thus had small standard deviations. We tested different thresholds, including zero, and none of them changed the results we present in this paper.

To calculate percent fluorescent change in a cell, the mean ΔF/F was calculated by projecting the ΔF/F matrix (time-point samples x pixels in the cell) onto the spatial filter a (pixels x1), and normalizing by the sum of the coefficients a_i_ to produce a weighted mean. The filter *a* was optimized from the deconvolution algorithm.

Around each cell, a local neuropil patch was defined by selecting an annulus with radii of 7 to 15μm centered around the middle of the cell body. Unless explicitly specified, all neuropil results shown in the present study were drawn from this neuropil patch size. We chose the minimum inner radius as 7 μm to minimize contamination from the cell signal onto the neuropil patch. In accordance with a previous report for the size of cell somata in mouse visual cortex [7], this was indeed large enough to exclude any pixels from other cell somata, by visual inspection. Other cell bodies, glia, and dark blood vessel regions were similarly excluded. After the application of HPF (cut-off at 0.05 Hz), the ΔF/F was calculated in single pixels with the same method as for the cell somata, and then averaged across all pixels within the patch.

The mean response to the visual stimulus was calculated for each ROI (cell or neuropil) after subtracting the mean ΔF/F response over the last 3 frames prior to stimulus onset. An aggregate response was computed by averaging the calcium signal over a period corresponding to the duration of the stimulus presentation (600ms), centered at the peak frame of the mean response computed across the cell population for stimuli at 100% and 40% contrast. This strategy was particularly important for anesthetized animals because the time course of the calcium signal was more prolonged [24], peaking near the end of the stimulus presentation. Cells were called visually responsive if they had a significant mean response across all stimuli presented at 100% contrast (i.e., ΔF/F > 0.5%). In the analysis that follows we used visually response cells and their surround neuropil patches.

Fano factors were calculated from ΔF/F (%) responses individually for the 6 stimulus conditions, and averaged across the two grating orientations presented for each given contrast. Mean Fano factors of cells and neuropil patches, and their ratios (Fcell/Fneuropil), were calculated across each FOV. The overall mean and SEM of the ratio is reported across FOV’s. Note that even though Fano factors were calculated here using ΔF/F (%) responses instead of spike rates, they are still useful for comparing the relative variability between cell and neuropil and across different brain states.

### Noise Correlation Analysis

For noise correlation analysis, pairwise Pearson’s correlation coefficients between pairs of cells, pairs of neuropil-patches and between cells and neuropil-patches, were calculated for each stimulus condition. The noise correlation coefficients were calculated by considering all individual trials within each stimulus condition. Then, noise correlation coefficient values of two directions in each contrast were averaged to obtain a mean noise correlation coefficient at the given contrast. For cells, estimated spike responses (see above) were used to calculate noise correlation coefficients. Neuropil-to-neuropil and cell-to-cell noise correlations were grouped separately according to their distance and averaged to yield the mean noise correlation coefficient as a function of distance. Linear fits to these plots were obtained per FOV and used to compare between the linear decays of neuropil-to-neuropil and cell-to-cell noise correlations by performing a statistical test for difference between the two slopes [25] as follows:

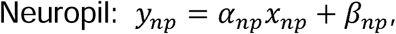

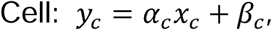
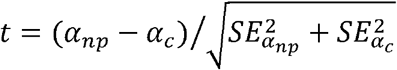, where SE indicates the standard error of a slope.

*t* ~ *T*(*n_np_ + n_c_* − 4), where T is the t-distribution, and *n_np_ and n_c_* are sample numbers of neuropil and cell data, respectively. This slope test was also performed between brain states within either cell or neuropil noise correlation.

We also measured *noise* cross-correlations between the cell soma and a set of adjacent, 2 μm-thick neuropil annuli located at progressively larger distances from the center of the cell soma. Briefly, 1) 2 μm-thick annular neuropil patches whose radii increased incrementally (31-33 μm, 51-53 μm, and so forth) were defined, 2) the mean cross-correlations between the cell soma and its corresponding neuropil patches were measured across each FOV, and 3) the overall mean and standard error of the mean (SEM) were calculated across all FOV’s. To avoid local neuropil contamination of cell somata, we restricted the correlation analysis to cell and neuropil patches that were at least 30 μm apart from each other.

#### Neuropil correlation analysis after linear subtraction of adjacent cell response

To estimate the contribution of single cells’ responses to neuropil activity, single cells’ responses weighted with a common scale value were subtracted from neuropil responses. The common scale value was estimated through a linear regression model: *y* = *Χβ*, where *y* is an nx1 vector composed of n neuropil patch responses, *X* is a nx2 vector composed of an nx1 vector of cell responses and an nx1 vector of 1s, *β* is a 2×1 vector, composed of a scale value and a bias. After subtracting cell responses with the common scale value *β*, the residual neuropil responses, *y_r_* = *y* − *Χβ*, were used for correlation analysis to measure the linear subtraction effects. Even though the use of a common scale value across all pairs of cells and adjacent neuropil patches within an FOV does not take into account varying interactions of individual pairs, the overall effects from the linear subtraction can still be assessed under the assumption that overall effects are similar across FOV’s and brain states.

### Estimating Decoding Accuracy of Cells and Neuropil Patches for Direction Discrimination

To compare the decoding accuracy for direction discrimination between cells and neuropil patches, optimal linear decoding was used [26]. First, the responses of visually responsive cells to the stimulus were transformed into a vector representing a population rate code in each trial. Then, 10-fold cross-validation tests were performed by leaving out 10% of the data for testing, and training the classifier with the remaining 90%. Discriminability between vertical and horizontal gratings was measured separately for each contrast level.

Decoding accuracies of n-element populations were calculated by selecting *n* cells or *n* neuropil patches randomly from each FOV in each trial, collecting their responses into a population vector, calculating the vector’s direction decoding accuracy, and averaging decoding accuracies across vectors drawn independently across 1000 trials. The decoding accuracy in each trial was calculated by averaging 10 decoding accuracy values from 10-fold cross-validation tests. Then, the overall mean and SEM were obtained across FOVs. In this procedure, a smaller neuropil patch size (annulus radii of 7 to 11μm) was chosen, which reduced the overlap of pair-wise neuropil patches to <0.5% of the patch area to minimize artificially generated signal correlations.

Neuropil decoding accuracies were also compared across a range of patch sizes: 7-11 μm, 7-50 μm, 7-100 μm, …, 7-250 μm radii. Each patch was centered around the corresponding cell, and thus the overlap ratio between patches increased with patch size. As neuropil patch size increased, some neuropil patches completely overlapped with other patches (i.e., <0.5% for 7-11 μm to >80% for 7-250μm). Therefore, as optimal linear decoders require full ranks of data, we removed some patches when adding a new patch in the population vector did not increase the rank of data. Again, all decoding accuracies were computed from 10-fold cross-validation tests.

#### Direction discriminability within single cells and neuropil-patches

To assess direction discriminability of cell and neuropil activity, we used d’:

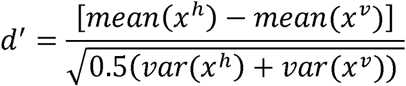

where *x* represents a trial-response of either cell or neuropil-patch conditioned on stimulus direction, which is denoted with superscript *v* (vertical grating) and *h* (horizontal grating).

Then, we compared the spatial organization between the d’s of cells versus their local neuropil-patches by calculating a spatial Pearson correlation coefficient between the corresponding *d’* values, within each FOV. To assess the statistical significance of the spatial correlation, we built a null distribution from correlation coefficient values obtained from 10,000 shuffled dataset. Each shuffled surrogate dataset was generated by randomly shuffling the spatial order of cells, thereby destroying the spatial relationship between cells and neuropil patches. From the null distribution, the statistical significance of the original spatial correlation between cell and neuropil-patch d’ values could be assessed.

## Results

### Visually-Evoked Neuropil Response Strength and Response Reliability

To explore visually evoked neuropil responses, we measured and compared percent fluorescence change (ΔF/F) seen in neuropil-patches versus cell somata. Both neuropil and cell (hereafter, the word cell refers to the soma of a neuron) responses were strongly modulated as a function of contrast, cell responses being generally weaker than neuropil responses (Figure 1D). Furthermore, the ratio of cell to neuropil-patch response strength decreased with falling contrast (Figure 1E), because somatic activity dropped faster than neuropil activity as contrast was lowered. This was true both in the awake state (AW) and under anesthesia (AN) (Figure 1E). Assuming that neuropil activity reflects aggregate inputs to the cell, this suggests that the cell’s ability to “transform” integrated inputs into action potentials weakens at low contrasts.

As expected, changes in brain state affect both neuropil and somatic responses. Specifically, light (0.7% isoflurane) anesthesia markedly increased visually driven activity in the neuropil (Figure 1D) and less so in the L2/3 neurons themselves. As a result, the ratio of visual response strength in L2/3 cells versus adjacent neuropil patches was lower under light anesthesia than in the quiet awake state (Figure 1E). This is consistent with the interpretation that the L2/3 cells ability to “transform” their inputs into action potentials weakens with anesthesia.

### Response Reliability

We compared the reliability of cell responses to neuropil-patch responses when exposed to the repeated presentation of identical moving grating stimuli. Cell responses were highly variable whereas neuropil responses, even for small patches, showed much greater reliability. We calculated and compared the Fano factors of cells versus neuropil-patches using the calcium signal (variance/mean calculated using the ΔF/F (%) response; see methods). The Fano factors of neuropil annuli (radii 7-15 μm from the cell soma) were 5-20 times smaller than the Fano factor of their corresponding cell (P<1e-8; the main effect between cell versus neuropil-patch in two-way ANOVA; Figure 3A), reflecting the high reliability of aggregate neuropil responses. This suggests that the high randomness of cell firing results either i) from the cells’ own internal processes, or by ii) sub-selecting a specific subgroup of inputs with higher variability than seen in the aggregate neuropil activity. Naturally, Fano factors increased as the number of pixels in the neuropil patch decreased, but even small 5-pixel neuropil patches (~11 μm^2^; covering ~ 4-7 times smaller area than the cell body) still yielded ~2 times larger Fano factors than those of cells (1.86; Figure 3B). Mean neuropil Fano factors approached the cellular Fano factors for small (1 pixel = ~2.25 μm^2^) neuropil patch sizes, but then rapidly dropped for patches of bigger sizes (Figure 3B).

**Figure 3.**
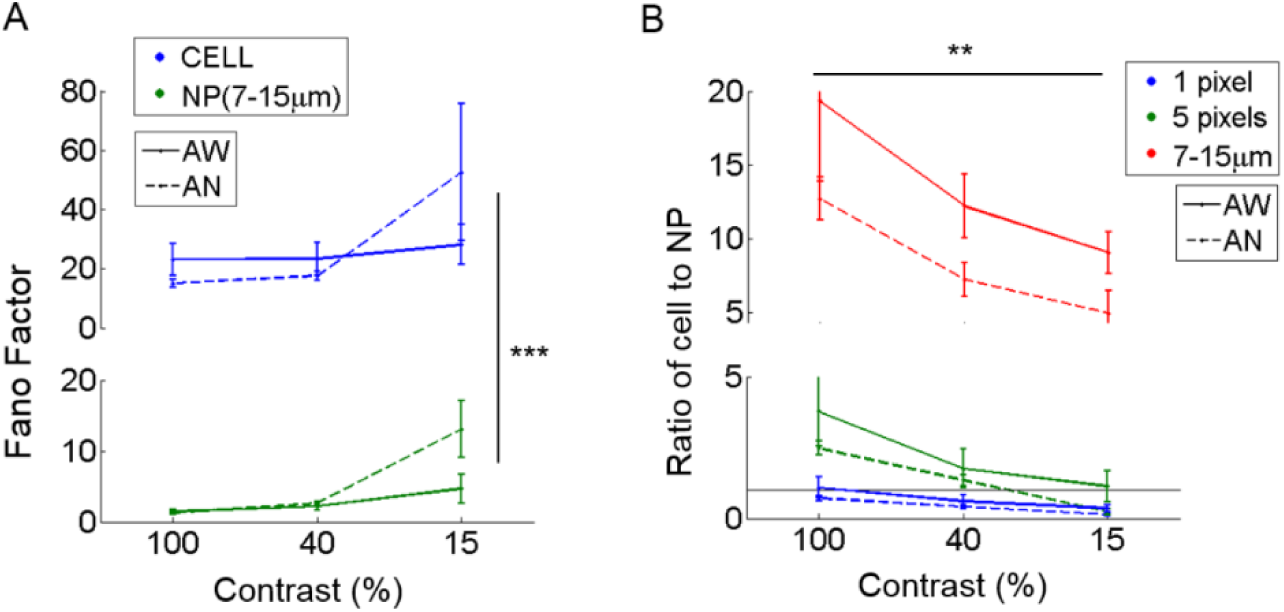
Fano factors of cells versus adjacent neuropil patches. Fano factors (variance/mean) were estimated in cells and nearby surrounding neuropil patches. Fano factors were calculated based on ΔF/F responses, and thus the absolute scale is different from that calculated using spike rates measured by electrophysiology. **(A)** Mean Fano factor across cells (CELL) versus adjacent annular neuropil patches (NP; radius 7-15 μm). Neuropil patches show very small Fano factors across all brain states and visual contrasts, meaning that neuropil responses to repetitions of identical stimuli are almost the same. In contrast, neurons have high Fano factors reflecting the large variability of neuronal responses. *** indicates a significance level of P<1e-8 in comparison between Fano factors of cells and neuropil-patches (Two-way ANOVA). **(B)** Ratio of the mean Fano factor across cells versus across neuropil patches. Neuropil patches of different sizes are illustrated in different colors. Neuropil patches with 1 or 5 pixels were defined by randomly selecting 1 or 5 pixels within annular neuropil patches centered around the cell extending from 7 to 10 μm. A ratio of 1 is shown as a black horizontal line. Mean neuropil Fano factors approach the cellular Fano factors for small (1 pixel = ~2.25 μm^2^) neuropil patch sizes, but then rapidly drop for patches of bigger sizes. ** indicates a significance level of P<5e-4 across contrasts (Two-way ANOVA).

The relative reliability of neuropil to cell responses varies as a function of visual contrast. Cell-to-neuropil Fano-factor ratios decreased with falling stimulus contrast in L2/3 (1 and 5 pixels, and annulus with 7-15μm radii; P<5e-4, in Two-way ANOVA; Figure 3B). This suggests that a fraction of the projections included in L2/3 neuropil patches fire more reliably than local L2/3 cells do as contrast increases. The relative reliability of cell to neuropil responses varied less strongly as a function of brain state. Typically cell-to-neuropil Fano-factor ratios were slightly higher in the quiet awake compared to the lightly anesthetized state (Figure 3B), however significance was not reached (P>0.5 for 1- and 5- pixels, and P=0.06 for annulus radii with 7-15μm in Wilcoxon rank-sum test), suggesting that changes in brain state may affect more uniformly the local and non-local neural processes that constitute the L2/3 neuropil patches examined.

In summary, we found that visually evoked responses of both cells and adjacent neuropil patches depend on brain state, and that the ratio of cell to neuropil response magnitude is much larger during quiet wakefulness than under light anesthesia. Neuropil patch responses to drifting gratings were always more reliable than L2/3 cell responses, setting a limit on the degree to which the spatially coherent signal carried by the neuropil, which may arise as a projection from other areas, can fluctuate randomly from trial to trial. Furthermore, neuropil activity does not simply reflect the linear sum of nearby cell activity since cell-to-neuropil fano-factor-ratios change as a function of visual contrast.

### Neuropil-to-Neuropil, Cell-to-Neuropil and Cell-to-Cell Noise Correlations

Noise correlation analysis is thought to provide information about the spatial organization of shared “internal” input modulations due to brain state fluctuations [9,10] and other global modulatory inputs [27] that influence neuronal activity. Noise correlations between different disjoint neuropil patches were high (~0.82 under light anesthesia vs ~0.67 in quiet wakefulness, P < 1e-8, Wilcoxon rank-sum test; Figure 4A, *Left*), in line with prior observations that demonstrated the high spatial coherence of the neuropil signal [2]. Notably, these values are strikingly different from the strength of cell-to-cell noise correlation coefficients, which were much lower (< 0.1 in anesthetized, ~0.05 in the awake state; Figure 4A, *Right*). One possible explanation for this difference is that a local volume of neuropil integrates numerous synaptic processes [6,7] that may extend over considerable cortical distances, providing the same signal to different neuropil patches. Alternatively, the aggregate activity of neuropil processes may contain a spatially coherent signal, which is filtered out in pairwise cellular noise correlations by the stochastic firing of individual units.

**Figure 4.**
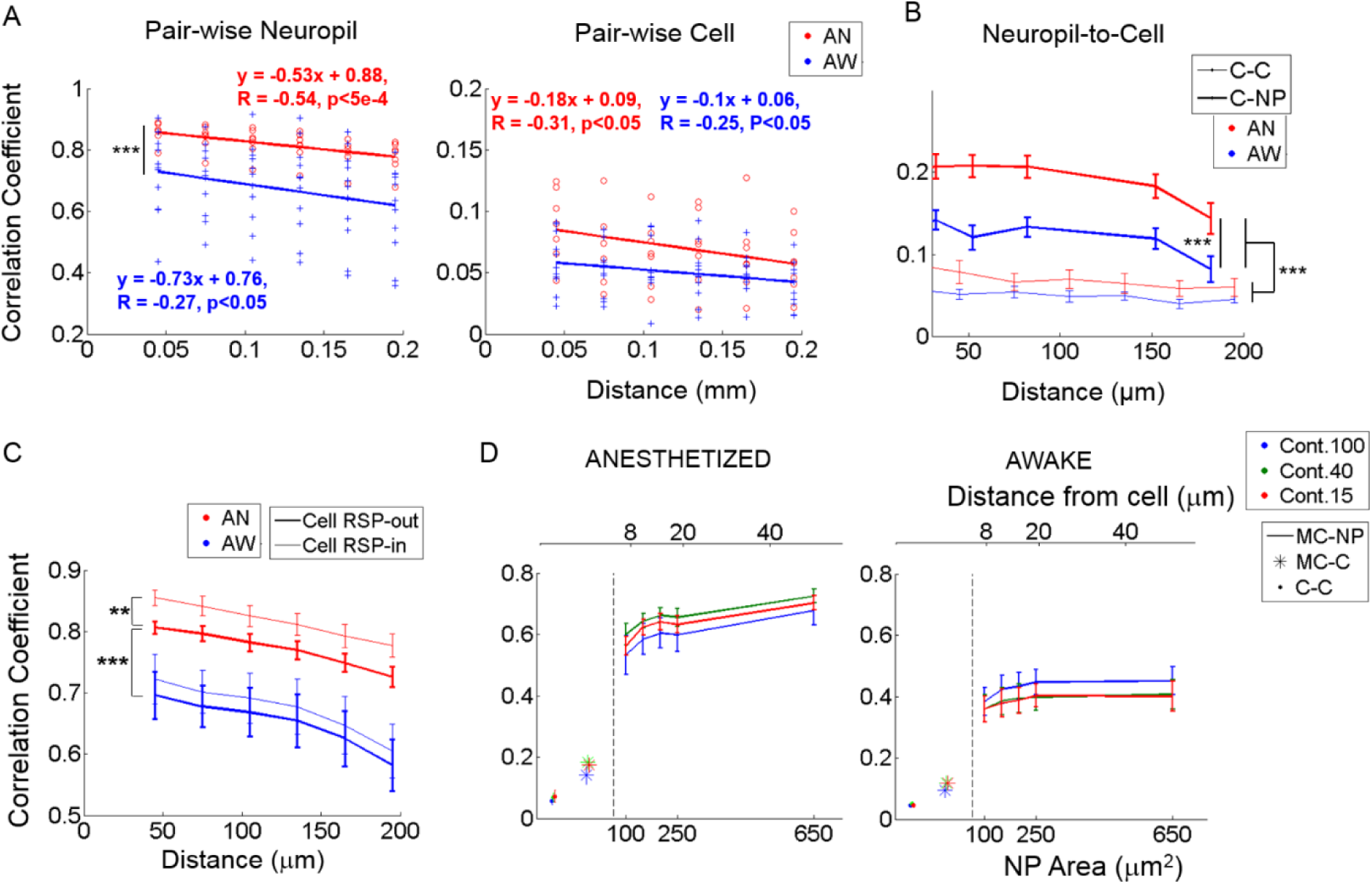
Noise correlations. **(A)** Noise correlation between neuropil-patches (*left*) and between cells (*right*) as a function of distance. In (A), each value marked with either a red circle (anesthetized) or a blue cross (awake) represent the mean correlation across all pairs belonging to the corresponding distance bin within an FOV. Bin size = 30 μm. While neuropil coefficients decay at a steeper slope than cell coefficients, (P<0.05, t-test for the two independent slopes), regardless of brain state, the relative (%) decrease over the same distance is ~ 3 times larger for cells (~30% for cells versus ~12% for neuropil patches). **(B)** Noise correlation between neuropil-patches and cells as a function of distance (bold line, C-NP). ‘C-C’ refers to pairwise cell-to-cell noise correlations. Error bars indicate mean ± SEM at distances used across FOVs. For (A-B), *** indicates P<1e-7 in Wilcoxon rank-sum test. **(C)** Linear contribution of single-cell response to neuropil-patch noise correlation. Mean noise correlations between neuropil patches were compared before and after subtracting corresponding single cells’ responses (see Methods). **, ***: P<5e-5, P<1e-8, Wilcoxon rank-sum test. ‘Cell RSP-in’: without linear subtraction. ‘Cell RSP-out’: after linear subtraction. Our basic observations remain unchanged after this correction. (D) Mean noise correlations between individual cells (C-C), between single cells and the mean response from all other cells except the cell itself (MC), and between the mean cell response (MC) and neuropil patches (NP) as a function of neuropil patch size and its distance from the centered cell. ‘X-Y’ represents noise correlation between ‘X’ and ‘Y’; for example, *MC-NP*: correlation between MC and NP. *AN*: Anesthesia, *AW*: Quiet wakefulness. *Cont*.: Visual contrast. Cell responses were measured from deconvolved spike rates. In plots, solid line (with error bar) represents the mean (with SEM) across FOV’s (n=7 and 11 from anesthetized and awake animals, respectively).

Despite the high spatial coherence of neuropil activity, neuropil-patch to neuropil-patch noise correlation strength decayed substantially as a function of cortical distance regardless of brain state (Figure 4A). For example, neuropil patches <40 μm apart from each other had significantly stronger noise correlations compared to patches located 180 μm to 210 μm from each other (9% and 15% decrease for the anesthetized, awake state respectively; p<0.05, *one-tailed Welch’s t-test*; Figure 4A, *Left*). This was true both in quiet wakefulness and under light anesthesia. In contrast to neuropil inter-patch noise correlations (Figure 4A, *Left*), pairwise cell noise correlations were flatter as a function of distance (slopes significantly different at P<0.05 for both brain states, t-test for two slopes; Figure 4A, *Right*), decreasing more slowly with increasing distances [13, 28–31]. Nonetheless, because they start at a much lower value, the relative decrease of cell-to-cell noise correlation strength was larger for the same cortical distance (i.e., <40 vs 180-210μm), decreasing by 27% and 33% for the anesthetized, awake state respectively (Figure 4A). This may potentially reflect a difference in the spatial organization of neuropil processes, largely reflecting synaptic inputs, versus the spatial organization of L2/3 cellular activity. Certainly, the large spatially coherent component that imparts to the neuropil its high noise correlation strength appears to be largely filtered out in L2/3 cell output activity.

To further explore how cell responses are related to neuropil responses, we measured *noise* cross-correlations between cells and a series of annular neuropil patches centered at increasing distances from the cell soma. Patches were chosen to be 2 μm-thick annuli located at progressively larger distances from the cell center. Neuropil-to-cell cross-correlation coefficients were compared to cell-to-cell cross-correlation coefficients. Pairwise cell-to-cell noise correlation coefficients were calculated using spike rates estimated from the calcium signal (see Methods and Figure 2). Neuropil-to-cell noise_correlations were markedly lower than neuropil-patch to neuropil-patch ones, but still larger than cell-to-cell noise correlations (P<1e-30, Wilcoxon rank-sum test; Figure 4B). This suggests that neurons efficiently filter out the major part of the highly coherent, spatially uniform, neuropil modulation component. We also found that coherence between cell and neuropil responses is brain state-dependent, yielding significantly lower noise correlation coefficients in the quiet awake state versus under light anesthesia (P<1e-7, Wilcoxon rank-sum test; Figure 4B). This suggests a more heterogeneous input/output activity structure in the quiet awake state.

In summary, neuropil activity exhibits much stronger spatial coherence than cell activity during the repeated presentation of identical stimuli, at least up to distances of ~200μm and likely higher. The relative decrease of noise correlation coefficient strength as a function of distance is smaller in neuropil than in cells, and both cell and neuropil noise correlation coefficients depend on brain state, being smaller in quiet wakefulness versus light anesthesia. These observations suggest that L2/3 neurons are capable of filtering out the main, highly coherent, spatially uniform, neuropil modulation component, and that neuropil activity exhibits a more heterogeneous structure in the quiet awake state.

### Contribution of Cell Responses to nearby Neuropil Activity

To investigate the influence of the activity of individual cells on nearby neuropil activity, we subtracted somatic responses weighted with a common scale value from the adjacent neuropil-patch responses (annulus from 7 to 15 μm centered around the cell). The scale value was simply estimated across all cell-neuropil-patch pairs using a linear regression model between a dependent variable (i.e., the neuropil response) and a regressor (i.e., the corresponding cell response, see Methods for details). Then, we recalculated the noise correlations between neuropil patches. This approach removed the mean linear contribution of individual cells to neuropil activity and allowed us to determine to what degree the spatial structure of neuropil activity described in the previous section could have its origin in the activity of adjacent cells, as might occur, for example, via the back-propagation of action potentials through the cells’ own dendritic branches [32].

We found that subtracting somatic responses significantly reduced neuropil inter-patch noise correlations only under anesthesia (0.046 vs. 0.023, P<5e-5 vs. P = 0.24, Wilcoxon rank-sum test), without significantly affecting their profile over distance (P<1e-8, Wilcoxon rank-sum test between the two brain-states; Figure 4C). This is consistent with the observation that single cells are more strongly synchronized with surrounding cell populations and neuropil under anesthesia than in the quiet awake state [10].

Figure 4B shows that neuropil-to-cell noise correlations were considerably stronger than mean cell-to-cell noise correlations, across all distances measured (e.g., 0.18 vs. 0.06 under anesthesia and 0.12 vs. 0.04 for the quiet awake state, respectively, at 150 μm). The high correlation values imply that neuropil patches carry a substantial signal that is correlated with cell activity, even for cells that are quite distant from the location of the neuropil patch (~150 μm). Although this might initially appear counterintuitive, it may have an obvious reason: neuropil responses reflect aggregate synaptic activity and are more correlated with mean cell population activity than with individual cell activity. Noise correlation coefficients between individual cell responses and mean population responses were also much higher than the average of all pairwise cell-to-cell noise correlation coefficients. The reason is that the firing of individual cells is stochastic, which disproportionately lowers pairwise cell-to-cell noise correlation coefficients. To test this reasoning, we measured 1) the correlation coefficient between the response of individual cells and the mean cell response (MC) from all other cells except the cell itself (MC-C), and 2) the correlation coefficient between MC and individual neuropil patch responses (MC-NP). As predicted, MC-C noise correlation coefficients were higher than mean pairwise cell-to-cell noise correlation coefficients, and MC-NP correlation coefficients were even higher (Figure 4D).

Noise correlations were affected by brain state, being generally higher during anesthesia (Figure 4D, compare left to right plot). This was particularly true for the MC-NP cross-correlation values, which were 80% higher under anesthesia (0.74 at 50 μm) versus in the quiet awake state (0.41 at 50 μm) (Figure 4D). It is therefore likely that under anesthesia, the population of cells is strongly driven by the activity of shared inputs. It is also likely that in the quiet awake state, sparse and more localized patterns of firing activity are predominant [33,34], leading to smaller noise correlation coefficients.

In summary, removing the calcium signal component that may contribute to the neuropil by cells does not change the spatial profile of the neuropil-to-neuropil noise correlation structure. Furthermore, cell to neuropil-patch noise correlations are significantly stronger than the mean cell-to-cell noise correlation. This is likely due to the stochastic nature of cell firing, which disproportionately lowers pairwise cell-to-cell noise correlation coefficients.

### Decoding Direction Information from Neuropil Patch Populations

Mice lack orientation columns and are thought to have a salt and pepper pattern of orientation/direction preference [35]. However, a recent study suggests that orientation preference may not be entirely randomly distributed across neurons, as neighboring cells show greater similarity in orientation preference [36]. Neuropil neighborhoods of appreciable size contain numerous cell processes. Assuming orientation/direction preference is randomly distributed across these processes, neuropil-patches surrounding neurons should show little, if any, orientation/direction bias. Alternatively, neuropil processes may be organized in regions with coherent orientation/direction preference, which changes gradually along the cortical surface. In what follows we ask whether neuropil activity conveys sufficient information to decode visual direction and how this compares to the direction discrimination accuracy of nearby cell populations (see Methods).

We find that populations of neuropil-patches show high decoding performance, commensurate with that of populations of neighboring neurons (Figure 5A, *Left*), both in quiet wakefulness and under light anesthesia. This high decoding performance from neuropil-patch populations did not result just from differences in the global modulation of aggregate neuropil activity depending on stimulus direction since direction selective information is essentially lost and discrimination accuracy drops to chance levels when we average over all neuropil patches within an FOV, (Figure 5A, *Right*).

**Figure 5.**
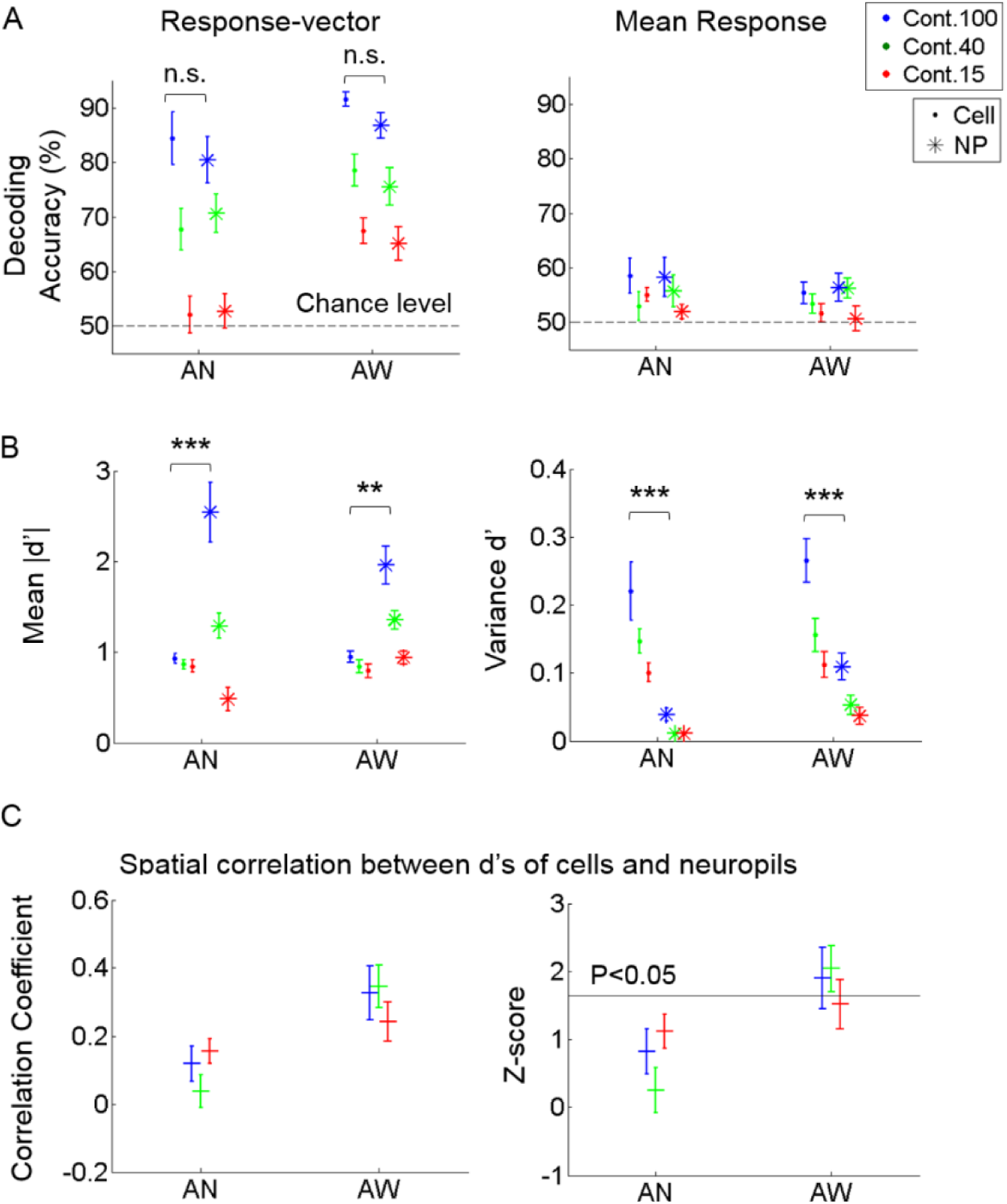
Decoding performance of cell versus neuropil response for stimulus direction. **(A)** Decoding accuracy from response vectors composed of cells versus neuropil-patches (*left*) and from the mean response across cells versus neuropil-patches within FOV’s (*right*). Decoding accuracy was not significantly different for neuropil-patch versus somatic response-vectors regardless of brain state (not significant in Two-way ANOVA). Response-averaging across population elements dropped the decoding performance to near chance level for both cells and neuropil patches. **(B)** Mean of absolute d’ values (*left*) and variance of d’ values (*right*) within each FOV, averaged across FOV’s. d’: discriminability between vertical versus horizontal grating conditions for single cell or neuropil-patches. On average, while the mean of absolute d’ values within a FOV is larger for neuropil-patches than for cells, the variance of d’ values is smaller across neuropil-patches. **, ***: P<5e-4, 1e-7 in Two-way ANOVA. (C) Spatial correlation between the d’s of cells versus their local neuropil-patches (*left*) and the corresponding z-score (*right; see methods*). Cell and neuropil patch *d’* vectors are significantly spatially correlated only in the quiet awake state (P<0.05). In all plots, ‘.’ stands for cell, ‘*’ for neuropil, ‘+’ represent the overall mean across FOVs, and error bars SEM across FOVs (n=7, 11 for anesthetized and awake animals). *AN*: Anesthesia, *AW*: Quiet wakefulness.

To better understand the sources of high neuropil decoding performance, we first assessed whether the spatial organization of neuropil patches for stimulus encoding is correlated with those of adjacent cells. To this end, we first calculated the extent of discriminability between the two stimulus directions, called d’, for single cells and neuropil-patches, respectively (see Methods). Then, we calculated a spatial Pearson correlation coefficient between the *d*’ values of cells and their local neuropil-patches within each FOV and compared it with a null distribution generated from surrogate data, where the cells’ spatial locations were randomly shuffled (see Methods). This test showed that neuropil activity was correlated with adjacent cell activity in terms of spatial organization relevant to stimulus encoding (Figure 5C). In particular, significance was only reached in the quiet awake state (Figure 5C, *Right*). Nonetheless, the magnitude of this correlation was not sufficient to explain the high neuropil decoding performance, particularly in the anesthetized state, in which the absolute correlation coefficient among d’ values was relatively weak and not significant (Figure 5C, *Left*). To further investigate the reason for the high decoding performance of neuropil activity, we measured the mean and variance of cell and neuropil-patch *d’* values within each FOV. On average, *d’* variances were significantly smaller, and mean absolute *d’* values were significantly larger across neuropil-patches compared to cells (Figure 5B). Therefore, in terms of the capacity for information encoding, neuropil activity shows higher and more spatially uniform discriminability between the chosen stimulus conditions than cell activity.

We also examined how direction discrimination accuracy varies as a function of population-vector size and as a function of neuropil patch size. As expected, we found that including more cells or neuropil-patches increased decoding accuracy for both cell and neuropil-patch populations (Figure 6A). Interestingly, cell and neuropil-patch populations showed similar accuracy in decoding the (coarse) grating drifting-direction difference we tested across all population sizes tested. Neuropil-population decoding accuracy decreased as neuropil patch size increased (Figure 6B). However, regardless of brain state, the decay was slow, and high direction discrimination accuracies (e.g. ~75% at 100% contrast) were still present for patches with radius as large as 250 μm. This suggests that, due to the high response reliability of the neuropil, even a small amount of spatial inhomogeneity is enough to accurately decode the direction difference employed.

**Figure 6.**
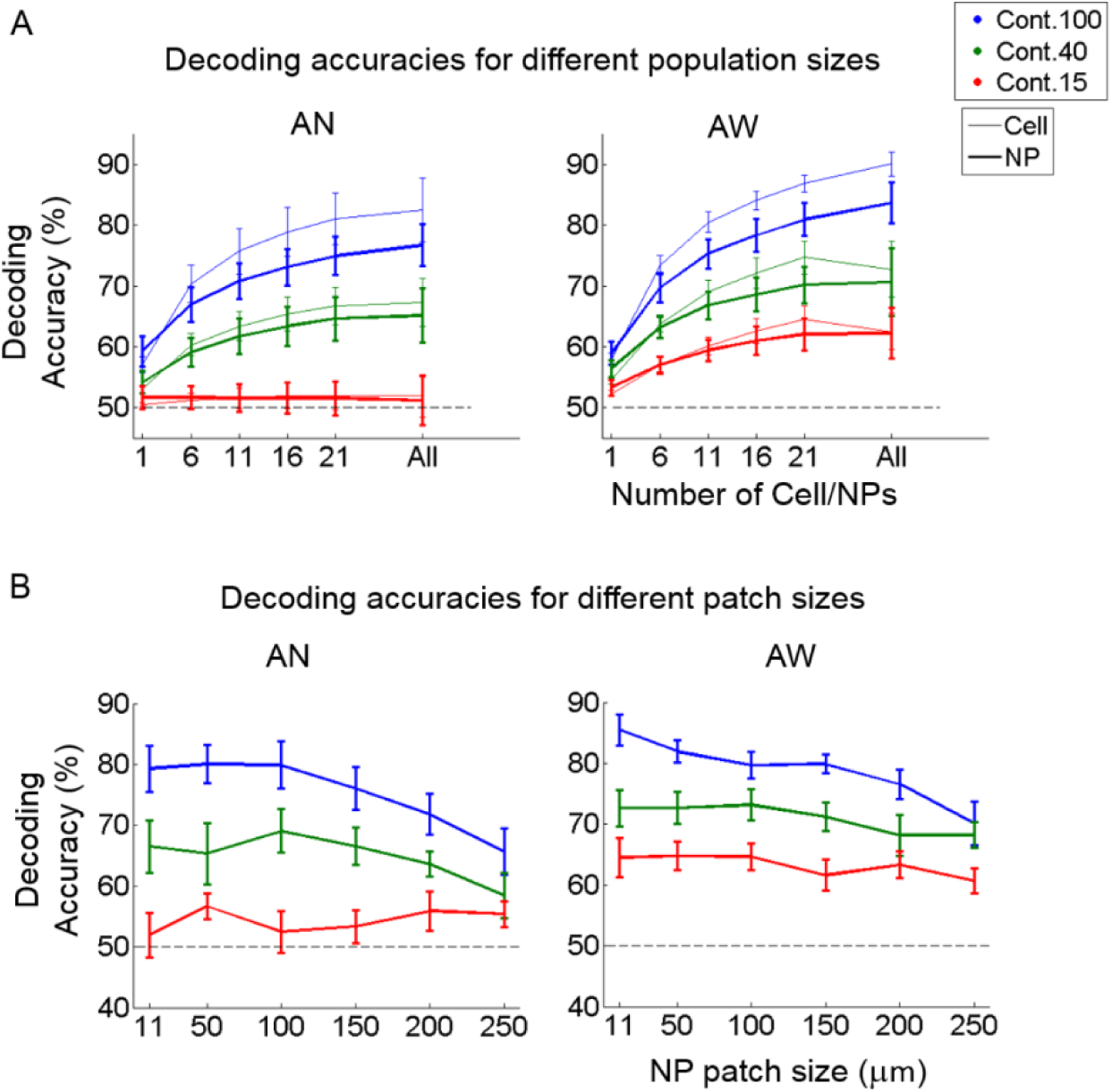
Decoding accuracy as a function of population-vector and neuropil-patch size. **(A)** Decoding accuracy as a function of population-vector size. Decoding accuracy of both cell and neuropil-patch populations increase in a similar way as a function of population-vector size. Decoding accuracies of the neuropil population are slightly lower for n>6, but this does not reach significance on the F-test. **(B)** Decoding accuracy as a function of neuropil patch size increases by enlarging the outer annulus radius, while keeping the inner radius constant at 7 μm. Note that decoding accuracy remains high well beyond neuropil patches with radius ~100 μm. Blue: 100%, green: 40%, red: 15%. Error bars represent SEM across FOV’s. ‘AN’: Anesthetized state. ‘AW’: Awake state. Dashed lines represent the chance level of accuracy.

## Discussion

We explored the properties of visually driven neuropil activity recorded in L2/3 of mouse area V1, how it relates to the activity of neighboring cells, and whether it can be effectively decoded to perform coarse stimulus orientation discrimination.

### Cell versus Neuropil Visual Response Strength and Reliability

The neuropil in layer 2/3 of mouse primary visual cortex (V1) showed strong visually evoked responses both under anesthesia and during quiet wakefulness. Neuropil ΔF/F responses were in fact stronger, on average, than cell ΔF/F responses at all contrasts (Figure 1). The ratio of neuronal to nearby neuropil ΔF/F responses depended on brain state. Specifically, this ratio was lower in the anesthetized compared to the awake state (Figure 1E) suggesting that synaptic processing is less efficient under anesthesia. Interestingly, the ratio of neuronal to neuropil response strength was higher for high contrast stimuli irrespective of brain state (Figures 1D, E). This suggests that at higher contrasts, the same aggregate neuropil activity input, as judged by the amplitude of neuropil calcium ΔF/F modulation, elicits higher levels of activity in target neurons.

Neuropil patch responses were always considerably more reliable than local L2/3 cell responses. This result may be thought of as largely expected, since a neuropil patch represents aggregate activity originating from the processes of many cells (see Figure 3). However, it is not altogether trivial, since it sets a limit on the degree to which the spatially coherent signal carried by the neuropil (which might arise as a result of input from other areas or subcortical structures) fluctuates randomly from trial to trial. The higher Fano factor seen in cell responses may be related to two factors: *1*) Higher randomness in cell firing may result from the cell’s own internal input/output processing as opposed to variability inherent to its inputs. For example, this elevated randomness in cell firing may be related with the mechanism called “iceberg effect”, which refers to increased variability in cellular firing that occurs when the firing threshold approaches the peak of stimulus-elicited membrane potential fluctuations[37]. *2*) Cells may sub-select a particular group of inputs, or difference of inputs, that displays higher variability.

For example, cell sub-threshold activity reflects the difference between excitatory and inhibitory inputs, and this difference, particularly after thresholding, might well be less reliable than the aggregate neuropil activity (which reflects the sum rather than the difference between inhibitory and excitatory inputs).

The Fano factors in both neurons and neuropil decrease at higher contrast (Figure 3A). However, the relative response variability of cells versus neuropil-patches, quantified by the Fano factor ratio, increases at higher contrast (see Figure 3B). The relative increase of cell response variability at higher contrasts may reflect contrast dependent variability in signal integration and spiking processes internal to the neuron (e.g., contrast-dependent iceberg effect [37]) or heterogeneous gain control modulation across different types of internal processes that provide input to the neuron. In contrast, a change of brain-state from quiet wakefulness to light anesthesia appears to have less influence over the cell-to-neuropil Fano factor ratio (no significant increase of the ratio in the quiet awake state compared to under light anesthesia; Figure 3). This suggests that brain-state changes explored here modulate the relative reliability of cell to neuropil visual responses less strongly than changes in visual contrast.

In summary, the neuropil shows strong visually evoked responses that depend both on brain state and visual contrast. Visual responses of neuropil patches with an area >11 μm^2^ (i. e. at least 1/5 the area of a cell soma) were considerably more reliable than cell responses. Assuming that local neuropil activity is a reasonable representation of the major inputs that a cell receives [2], this suggests that the change in neuronal response variability as a function of visual contrast may be mediated in part by stochasticity in the cell’s own internal input/output processing. Alternatively, cells may sub-select a particular group of inputs, or difference between inputs, that displays higher variability.

### Noise Correlations of Neuropil versus Cell Responses

Noise correlations reflect the co-modulation of responses by internal, common, inputs [10, 38–40]. Neuropil-to-neuropil noise correlations were overall very large (i.e., 0.6-0.85) even when neuropil patches were almost 200 μm apart from one another (see Figure 4A). In contrast, neuron-to-neuron pairwise noise correlations were much lower (see Figure 4B, C). The spatial profile of neuropil noise-correlated activity decayed linearly, dropping by ~11% by ~200 μm, approximately 3 times less than cell-to-cell noise correlated activity, which fell by ~30% (Figure 4A) over the same distance. Noise correlation profiles were brain-state dependent, showing higher correlation coefficients under light anesthesia versus quiet wakefulness, in agreement with [10]. Brain-state dependent changes in noise correlation strength were similar for noise correlations between neuropil-patches, pairs of neurons, or between neurons and neuropil-patches. Notably, these results remained valid even after subtracting the response profile of single cells from the response profile of nearby neuropil patches (Figure 4C), suggesting that local neuronal activity (or possibly signal contamination from neuronal somata) is not the main cause of our observations.

In agreement with [10,40], noise correlation strength between pairs of neurons was extremely low. However, neuropil-to-cell noise correlations were significantly higher than the mean pairwise cell correlation for a range of cortical distances from 50 to 150 μm regardless of brain state (Figure 4B). Kerr et al. [2], which used the same calcium dye as we used, have argued that neuropil modulations mostly reflect signals arising from presynaptic axonal activity. This suggests that the high spatial coherence in the neuropil reflects shared input across all embedded cells. This shared component is also captured by the mean activity across all somata (MC) in the FOV (Figure 4D). However, because individual cells fire stochastically, pair wise inter-neuronal correlations weaken, resulting in the following relationships: NP-MC > MC-C ~ NP-C >> C-C (Figure 4D). The fact that individual cells are strongly correlated to mean cell activity suggests that, at least up to distances of ~200 μm, they do receive a substantial amount of shared internal input. This finding is supported by recent studies showing that single cell activity is coupled to population activity [41]. In addition, the stronger correlation between single cell and mean cell activity found under light anesthesia (Figure 4) is consistent with prior results showing that stronger co-modulation occurs across cells during anesthesia versus in quiet wakefulness [10].

In summary: 1) Neuropil activity in mouse V1 is highly correlated over large cortical distances. 2) This large spatially coherent component is predominantly but not completely filtered out in L2/3 cell output activity (Figure 4). 3) This is reflected in the fact that single-cell to neuropil-patch noise correlation coefficient strength (though much lower than neuropil-patch to neuropil-patch noise correlation coefficient strength) remains flat as a function of distance across >150 μm (Figure 4B). 4) The stochasticity and sparseness of cell firing is likely responsible for the much lower cell-to-cell noise correlation coefficient strength (Figure 4B). 5) Both neuropil and cell noise correlation coefficients decrease in quiet wakefulness. Finally, 6) the spatial profile of neuropil noise correlations, which likely reflects primarily the coherence of L2/3 inputs, decays slowly over distance.

### Direction Discrimination from Neuropil Activity

Visually driven neuropil responses are both strong and reliable suggesting that they contain significant information about visual stimulus contrast. It is an open question whether they also contain significant information about stimulus orientation/direction. We found that populations of neuropil patches ranging in size from ~220 μm^2^ (radius ~8.4μm) to ~200,000 μm^2^ (radius ~250μm) discriminated moving grating direction of motion accurately, on par with corresponding cell populations (Figures 5–6).

The high neuropil decoding performance was not due to trivial contamination of the calcium signal by nearby cell activity because: 1) the choice of neuropil patch inner diameter (>7 μm radius from the cell center) carefully excluded the region of optical contamination in the X-Y plane, 2) there was, at best, a weak correlation in discriminating power (d’) between cells and nearby neuropil patches (Figure 5C), and 3) decoding performance remained high even for larger (>100 μm in radius; Figure 6B) patches, which aggregate multiple processes with “salt-and-pepper” direction organization [35]. Rather, the high decoding performance of neuropil activity was related to the high neuropil response reliability and corresponding increased sensitivity to spatially distributed information, i.e. high d’ values (Figure 5B).

A potential caveat is that we computed neuropil discrimination accuracies for two orthogonal directions, and it is not clear how the observed discrimination performance would translate to stimuli that differ by smaller orientation/direction angles. Nonetheless, we provided clear evidence that neuropil patches do encode sufficient direction information to distinguish between two coarse (90 degree difference) directions (Figure 5). Interestingly, the high decoding accuracy of neuropil-patch populations are maintained even if we select patches of relatively large size (>100 μm in radius), which reflect the aggregate activity of a multitude of neuronal processes (Figure 7B).

## Conclusions

Neuropil responses to visual stimuli (moving gratings) in layer 2/3 of mouse V1 are more reliable and more strongly modulated than somatic responses. Stimulus independent fluctuations in neuropil activity are strong and highly correlated across the cortical surface up to distances of at least 200 μm. This contrasts with cell-to-cell pairwise noise correlations, which are much weaker. Finally, despite the “salt & pepper” organization of orientation preference across V1 neurons [35], neuropil-patch populations show high accuracy for performing coarse direction discrimination, commensurate to the accuracy of corresponding cell populations. This remains true even for patches of radius >100μm. These observations underscore the dynamic nature and functional organization of the layer 2/3 neuropil signal and the underlying inputs it reflects.

## Acknowledgement

This work was supported by a Simons Foundation Pilot Award and R21 NS088457 from NINDS to SS. The authors thank Dimitri Yatsenko for help generating the bead sample.

---

**Supplementary Figure 1.**
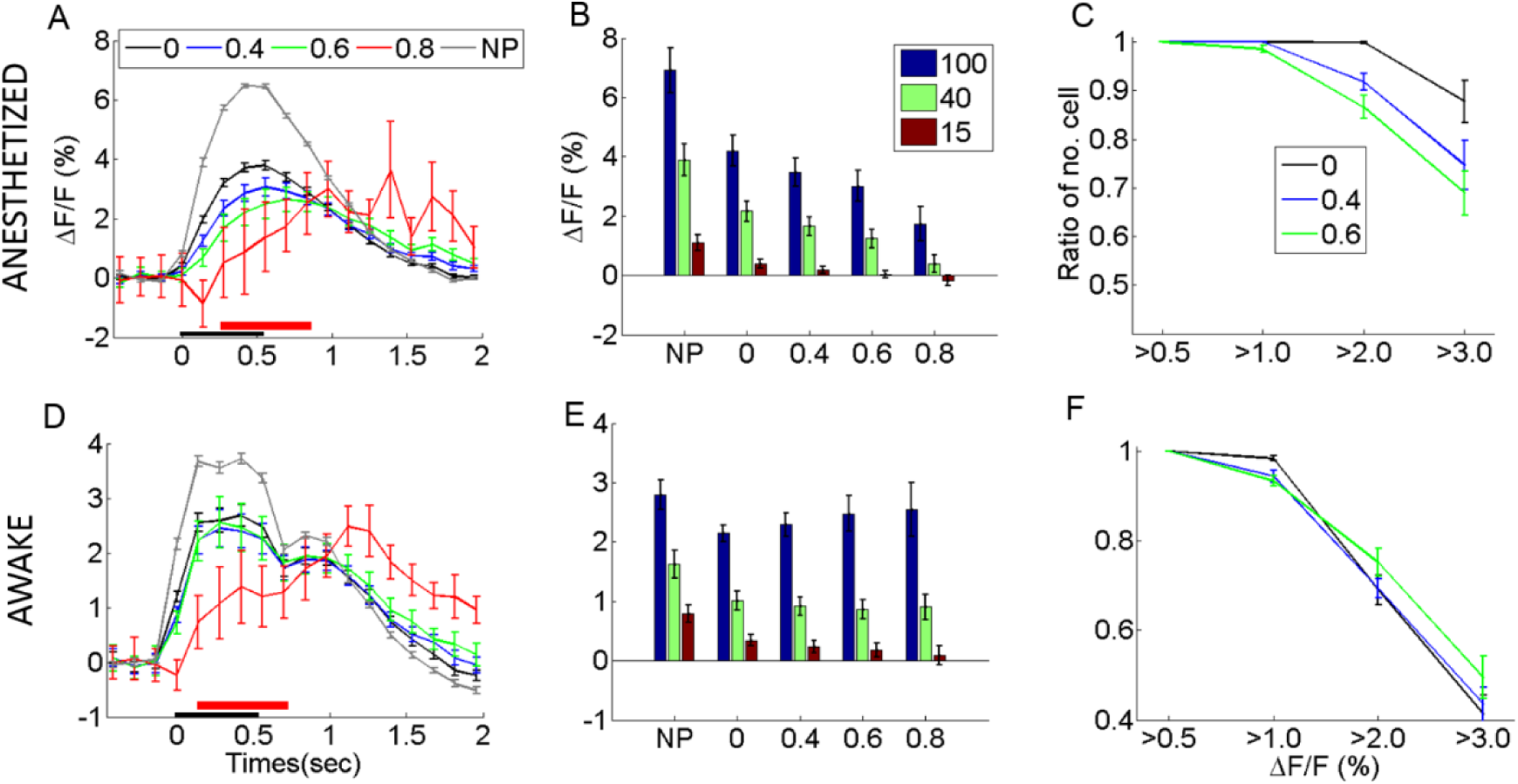
Effect of different levels of neuropil contamination correction on visual responses. Different levels of contamination correction (S) were used to correct the neuropil signal (see Methods), and the corrected ΔF/F response was plotted in the anesthetized (A - C) and awake states (D - F). (**A, D**) A typical example of the time course of the mean response to drifting gratings at 100% contrast, derived from the population of cells in a single field of view (FOV) in the anesthetized state (AN; top) versus the quiet wakefulness state (AW; bottom). The mean time course is corrected by subtracting the mean ΔF/F over 3 frames prior to stimulus onset. Cells shown had a mean response >0.5% to grating stimuli at 100% contrast. Curves with different color reflect different scale corrections (S ranging from 0 to 0.8). ‘NP’ indicates the neuropil signal from annular patches of radii of 7 to 15 μm. Error bars represent the standard error of the mean (S.E.M.). Black and red horizontal bars underlying the time courses correspond to the stimulus-on period and the time period for calculating the mean evoked response of ΔF/F, respectively. Image frame duration was ~130 msec (~7 Hz). The next frame following the onset of the stimulus was taken as t = 0. This means that the first frame at t = 0 sec has a timing jitter of 0-130 msec with respect to the stimulus onset, and thus it is not surprising that sometimes a small evoked response may be seen at t = 0 sec. Note that little change occurs in visually evoked responses for reasonable levels of contamination, i.e. S≤0.6. At higher levels of contamination, however, the cellular response is significantly affected by the correction suggesting that neuropil and cell responses cannot be separated at that stage. (B, E) Mean evoked responses to stimuli at 100%, 40%, and 15% contrast, derived from visually responsive cells in 7 AN and 11 AW FOV’s. Cells averaged had ΔF/F responses >0.5% at 100% contrast. ‘NP’ indicates the mean neuropil response. Here, the mean response of ΔF/F is corrected by subtracting the mean ΔF/F of 3 frames (~400 msec) prior to the stimulus onset, and thus slightly negative ΔF/F responses can exist. Note that for contrasts lower than 100% the neuropil contamination correction has a more significant effect. (C, F) Number of cells whose mean ΔF/F response to 100% contrast was greater than 0.5, 1.0, 2.0, 3.0% respectively, plotted for S = 0 (black), 0.4 (blue), and 0.6 (green). These numbers are normalized by the total number of cells with a mean response >0.5% under the given correction factor. Note that, in the anesthetized state, the neuropil contamination correction clearly reduces the fraction of cells with high response magnitudes, whereas in the awake state, where responses are overall weaker, this decline does not happen.

**Supplementary Figure 2.**
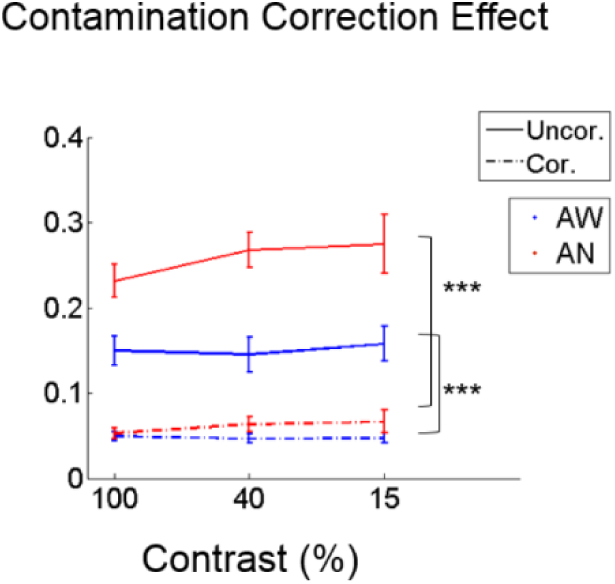
Decrease of noise correlations after neuropil contamination correction. Noise correlations of pairs of cells decrease after correcting for neuropil contamination with S = 0.6. ‘UnC.’ and ‘C.’ represent no correction of neuropil contamination (S = 0) and correction of neuropil contamination with S = 0.6, respectively. *** indicates a significance level of 1e-5.

## References

1. Grewe BF, Langer D, Kasper H, Kampa BM, Helmchen F (2010) High-speed in vivo calcium imaging reveals neuronal network activity with near-millisecond precision. Nat Methods 7: 399–405.

2. Kerr JN, Greenberg D, Helmchen F (2005) Imaging input and output of neocortical networks in vivo. Proc Natl Acad Sci U S A 102: 14063–14068.

3. Gobel W, Helmchen F (2007) In vivo calcium imaging of neural network function. Physiology (Bethesda) 22: 358–365.

4. Kerlin AM, Andermann ML, Berezovskii VK, Reid RC (2010) Broadly tuned response properties of diverse inhibitory neuron subtypes in mouse visual cortex. Neuron 67: 858–871.

5. Bonin V, Histed MH, Yurgenson S, Reid RC Local diversity and fine-scale organization of receptive fields in mouse visual cortex. J Neurosci 31: 18506–18521.

6. Chklovskii DB, Schikorski T, Stevens CF (2002) Wiring optimization in cortical circuits. Neuron 34: 341–347.

7. Braitenberg V, Shchuz A (1998) Cortex: Statistics and Geometry of Neuronal Connectivity. Berlin: Springer-Verlag.

8. Campagna JA, Miller KW, Forman SA (2003) Mechanisms of actions of inhaled anesthetics. N Engl J Med 348: 2110–2124.

9. Niell CM, Stryker MP (2010) Modulation of visual responses by behavioral state in mouse visual cortex. Neuron 65: 472–479.

10. Ecker AS, Berens P, Cotton RJ, Subramaniyan M, Denfield GH, et al. (2014) State dependence of noise correlations in macaque primary visual cortex. Neuron 82: 235–248.

11. Nimmerjahn A, Kirchhoff F, Kerr JN, Helmchen F (2004) Sulforhodamine 101 as a specific marker of astroglia in the neocortex in vivo. Nat Methods 1: 31–37.

12. Margrie TW, Meyer AH, Caputi A, Monyer H, Hasan MT, et al. (2003) Targeted whole-cell recordings in the mammalian brain in vivo. Neuron 39: 911–918.

13. Golshani P, Goncalves JT, Khoshkhoo S, Mostany R, Smirnakis S, et al. (2009) Internally mediated developmental desynchronization of neocortical network activity. J Neurosci 29: 10890–10899.

14. Brainard DH (1997) The Psychophysics Toolbox. Spat Vis 10: 433–436.

15. Michelson A (1927) Studies in Optics: U. of Chicago Press.

16. Guizar-Sicairos M, Thurman ST, Fienup JR (2008) Efficient subpixel image registration algorithms. Opt Lett 33: 156–158.

17. Chen TW, Wardill TJ, Sun Y, Pulver SR, Renninger SL, et al. (2013) Ultrasensitive fluorescent proteins for imaging neuronal activity. Nature 499: 295–300.

18. Vogelstein JT, Packer AM, Machado TA, Sippy T, Babadi B, et al. (2010) Fast nonnegative deconvolution for spike train inference from population calcium imaging. J Neurophysiol 104: 3691–3704.

19. Dempster AP, Laird NM, Rubin DB (1997) Maximum Likelihood from Incomplete Data via the EM Algorithm. Journal of the Royal Statistical Society Series B 39: 1–38.

20. Boyd SP, Vandenberghe L (2004) Convex optimization. Cambridge, UK; New York: Cambridge University Press, xiii, 716 p. p.

21. Kim S-J, Koh K, Lustig M, Boyd S, Gorinevsky D (2007) An Interior-Point Method for Large-Scale l1-Regularized Least Squares. IEEE JOURNAL OF SELECTED TOPICS IN SIGNAL PROCESSING 1: 606–617.

22. Kim H, Park H (2007) Sparse non-negative matrix factorizations via alternating non-negativity-constrained least squares for microarray data analysis. Bioinformatics 23: 1495–1502.

23. Grippo L, Sciandrone M (2000) On the convergence of the block nonlinear Gauss-Seidel method under convex constraints. Operations Research Letters 26: 127–136.

24. Haider B, Hausser M, Carandini M (2013) Inhibition dominates sensory responses in the awake cortex. Nature 493: 97–100.

25. Cohen J (1975) Applied multiple regression/correlation analysis for the behavioral sciences / Jacob Cohen, Patricia Cohen; Cohen P, editor. Hillsdale, N.J.: New York: Lawrence Erlbaum Associates ; distributed by Halsted Press Division of John Wiley.

26. Duda RO, Hart PE, Stork DG (2001) Pattern classification. New York: Wiley. xx, 654 p. p.

27. Polack PO, Friedman J, Golshani P (2013) Cellular mechanisms of brain state-dependent gain modulation in visual cortex. Nat Neurosci 16: 1331–1339.

28. Rothschild G, Nelken I, Mizrahi A (2010) Functional organization and population dynamics in the mouse primary auditory cortex. Nat Neurosci 13: 353–360.

29. Kerr JN, de Kock CP, Greenberg DS, Bruno RM, Sakmann B, et al. (2007) Spatial organization of neuronal population responses in layer 2/3 of rat barrel cortex. J Neurosci 27: 13316–13328.

30. Smith MA, Kohn A (2008) Spatial and temporal scales of neuronal correlation in primary visual cortex. J Neurosci 28: 12591–12603.

31. Denman DJ, Contreras D (2013) The structure of pairwise correlation in mouse primary visual cortex reveals functional organization in the absence of an orientation map. Cereb Cortex 24: 27072720.

32. Waters J, Larkum M, Sakmann B, Helmchen F (2003) Supralinear Ca2+ influx into dendritic tufts of layer 2/3 neocortical pyramidal neurons in vitro and in vivo. J Neurosci 23: 8558–8567.

33. Greenberg DS, Houweling AR, Kerr JN (2008) Population imaging of ongoing neuronal activity in the visual cortex of awake rats. Nat Neurosci 11: 749–751.

34. Sawinski J, Wallace DJ, Greenberg DS, Grossmann S, Denk W, et al. (2009) Visually evoked activity in cortical cells imaged in freely moving animals. Proc Natl Acad Sci U S A 106: 19557–19562.

35. Ohki K, Chung S, Ch'ng YH, Kara P, Reid RC (2005) Functional imaging with cellular resolution reveals precise micro-architecture in visual cortex. Nature 433: 597–603.

36. Ringach DL, Mineault PJ, Tring E, Olivas ND, Garcia-Junco-Clemente P, et al. (2016) Spatial clustering of tuning in mouse primary visual cortex. Nat Commun 7: 12270.

37. Priebe NJ, Ferster D (2008) Inhibition, spike threshold, and stimulus selectivity in primary visual cortex. Neuron 57: 482–497.

38. Bryant HL Jr., Marcos AR, Segundo JP (1973) Correlations of neuronal spike discharges produced by monosynaptic connections and by common inputs. J Neurophysiol 36: 205–225.

39. Shadlen MN, Newsome WT (1998) The variable discharge of cortical neurons: implications for connectivity, computation, and information coding. J Neurosci 18: 3870–3896.

40. Ecker AS, Berens P, Keliris GA, Bethge M, Logothetis NK, et al. (2010) Decorrelated neuronal firing in cortical microcircuits. Science 327: 584–587.

41. Okun M, Steinmetz NA, Cossell L, lacaruso MF, Ko H, et al. (2015) Diverse coupling of neurons to populations in sensory cortex. Nature 521: 511–515.

